# Temporal proteomic profiling reveals insight into critical developmental processes and temperature-influenced physiological response differences in a bivalve mollusc

**DOI:** 10.1101/2020.06.05.137059

**Authors:** Shelly A. Trigg, Kaitlyn R. Mitchell, Rhonda Elliott Thompson, Benoit Eudeline, Brent Vadopalas, Emma B. Timmins-Schiffman, Steven B. Roberts

## Abstract

**Background:** Protein expression patterns underlie physiological processes and phenotypic differences including those occurring during early development. The Pacific oyster (*Crassostrea gigas*) undergoes a major phenotypic change in early development from free-swimming larval form to sessile benthic dweller while proliferating in environments with broad temperature ranges. Despite the economic and ecological importance of the species, physiological processes occurring throughout metamorphosis and the impact of temperature on these processes have not yet been mapped out.

**Results:** Towards this, we comprehensively characterized protein abundance patterns for 7978 proteins throughout metamorphosis in the Pacific oyster at different temperature regimes. We used a multi-statistical approach including principal component analysis, ANOVA-simultaneous component analysis, and hierarchical clustering coupled with functional enrichment analysis to characterize these data. We identified distinct sets of proteins with time-dependent abundances generally not affected by temperature. Over 12 days, adhesion and calcification related proteins acutely decreased, organogenesis and extracellular matrix related proteins gradually decreased, proteins related to signaling showed sinusoidal abundance patterns, and proteins related to metabolic and growth processes gradually increased. Contrastingly, different sets of proteins showed temperature-dependent abundance patterns with proteins related to immune response showing lower abundance and catabolic pro-growth processes showing higher abundance in animals reared at 29°C relative to 23°C.

**Conclusion:** Although time was a stronger driver than temperature of metamorphic proteome changes, temperature-induced proteome differences led to pro-growth physiology corresponding to larger oyster size at 29°C, and to altered specific metamorphic processes and possible pathogen presence at 23°C. These findings offer high resolution insight into why oysters may experience high mortality rates during this life transition in both field and culture settings. The proteome resource generated by this study provides data-driven guidance for future work on developmental changes in molluscs. Furthermore, the analytical approach taken here provides a foundation for effective shotgun proteomic analyses across a variety of taxa.

## BACKGROUND

The Pacific oyster (*Crassostrea gigas*) is among the most ecologically and economically prominent bivalve molluscs given its contribution to biofiltration, habitat formation and stabilization, carbon and nitrogen sequestration, and international aquaculture revenue (~ $1.25 billion annually [1]). From a developmental perspective, it is a fascinating organism as it undergoes a complex transformation from a free-swimming planktonic larva to a sessile benthic juvenile. This involves two processes: settlement and metamorphosis. Once oysters acquire the ability to initiate and undergo morphogenesis (become competent), settlement commences typically 24 to 48 hours later where larvae drop out of the water column to the benthos, and use a newly developed foot to find appropriate substrate and secrete adhesive to attach to the substrate [2,3]. Then, metamorphosis typically occurs within a few to 72 hours and involves a complete rearrangement of organs, loss of larval organs including the velum and the foot, and development of new organs including gill-like ctenidia [4,5]. Complex physiology underlies both of these processes, involving neuroendocrine and immune functions and tightly-controlled gene expression programs, which is still not fully understood [6].

In addition to the physiological complexities of settlement and metamorphosis, the Pacific oyster is sensitive to abiotic and biotic factors during this particular life stage with substantial mortality occurring in both field and culture settings [4]. Past studies found that increased rearing temperature positively influenced survival with Pacific oyster larvae reared at 23°C leading to optimal recruitment success [7–9], and established 23°C as the standard aquaculture industry rearing temperature [10]. Yet hatcheries still frequently observe stochastic high mortality during this life stage. *C. gigas* has been observed to tolerate temperatures up to 37°C with 100% survival [11], and their extremely eurythermic trait has enabled their global distribution and culturing on nearly every continent including equatorial islands [12]. Particularly larvae and juveniles have been observed to tolerate temperatures up to 32°C when food is not limiting, with maximum growth rates and survival during settlement occurring above 27°C [13–16]. To better understand how temperature influences critical developmental processes and phenotype throughout metamorphosis, a comprehensive characterization of developmental physiological processes that occur throughout this life stage is needed.

Proteomics, a survey of the collection of all proteins and their abundances at a given time, is ideally suited to provide a basis for revealing the physiological complexities of development [17]. Specifically, untargeted shotgun proteomic profiling using liquid chromatography coupled tandem mass spectrometry (LC-MS/MS) can efficiently and accurately predict protein abundances by taking into account peptide count, spectral count and fragment-ion intensity [18]. LC-MS/MS has been effectively used to examine changes in biological processes during early larval development in bivalve molluscs prior to metamorphosis [19,20]. A two sample proteomic comparison of larvae just prior to metamorphosis with juveniles weeks after metamorphosis showed differences in proteins involved in tissue remodeling, signal transduction, and organ development, however the two proteomes had relatively low coverage (392 and 636 proteins, respectively) [21]. The proteomes of larvae just prior to metamorphosis held at 30°C showed changes that correlated with increased growth and calcification [22]. Nevertheless, deeper proteomic profiling of a broader sampling throughout metamorphosis and including temperature as a factor would bring more resolution to physiological processes occurring over time and those affected by temperature. Although untargeted large-scale multi-factor proteome studies pose the challenge of identifying core biological responses in the accompanying large complex datasets, applying multiple statistical approaches can reduce complexity of a large dataset to identify potential targets for diagnostics [23,24]. New analyses have been developed to consider temporal influences on multivariate datasets (e.g. analysis of variance simultaneous component analysis (ASCA)), and identify features impacted by specific experimental variables [25].

To comprehensively characterize developmental physiological processes that occur throughout settlement and metamorphosis in Pacific oyster domesticated in the Pacific Northwest region of the United States, we used LC-MS/MS to generate temporal proteomes from oyster larvae reared at two different temperature regimes to examine the basis for temperature-influenced phenotype differences. We developed an analysis framework that applies multiple statistical approaches to classify temporal and temperature-influenced developmental processes from proteomic responses. This longitudinal proteomic data paired with phenotypic data provide a high resolution perspective of the physiological mechanisms underlying metamorphic stages and how temperature influences them.

## RESULTS

### Larval performance and global proteome analysis

To comprehensively assess proteomes throughout metamorphosis we collected pools of whole animals at seven different time points starting at competency, the time when larvae have the ability to initiate and undergo metamorphosis (**Figure 1A**). After 6 days into the temperature treatment, we found minimal difference in the number of settled larvae between temperatures with 297,213 (29.7%) and 308,989 (30.9%) larvae settled at 23°C and at 29°C, respectively. However, larvae reared at 29°C tended to be larger in size at 24 days post-fertilization (dpf) (**Figure 1B**, **Additional File 1: Supplemental Table 1**).

**Figure 1.**
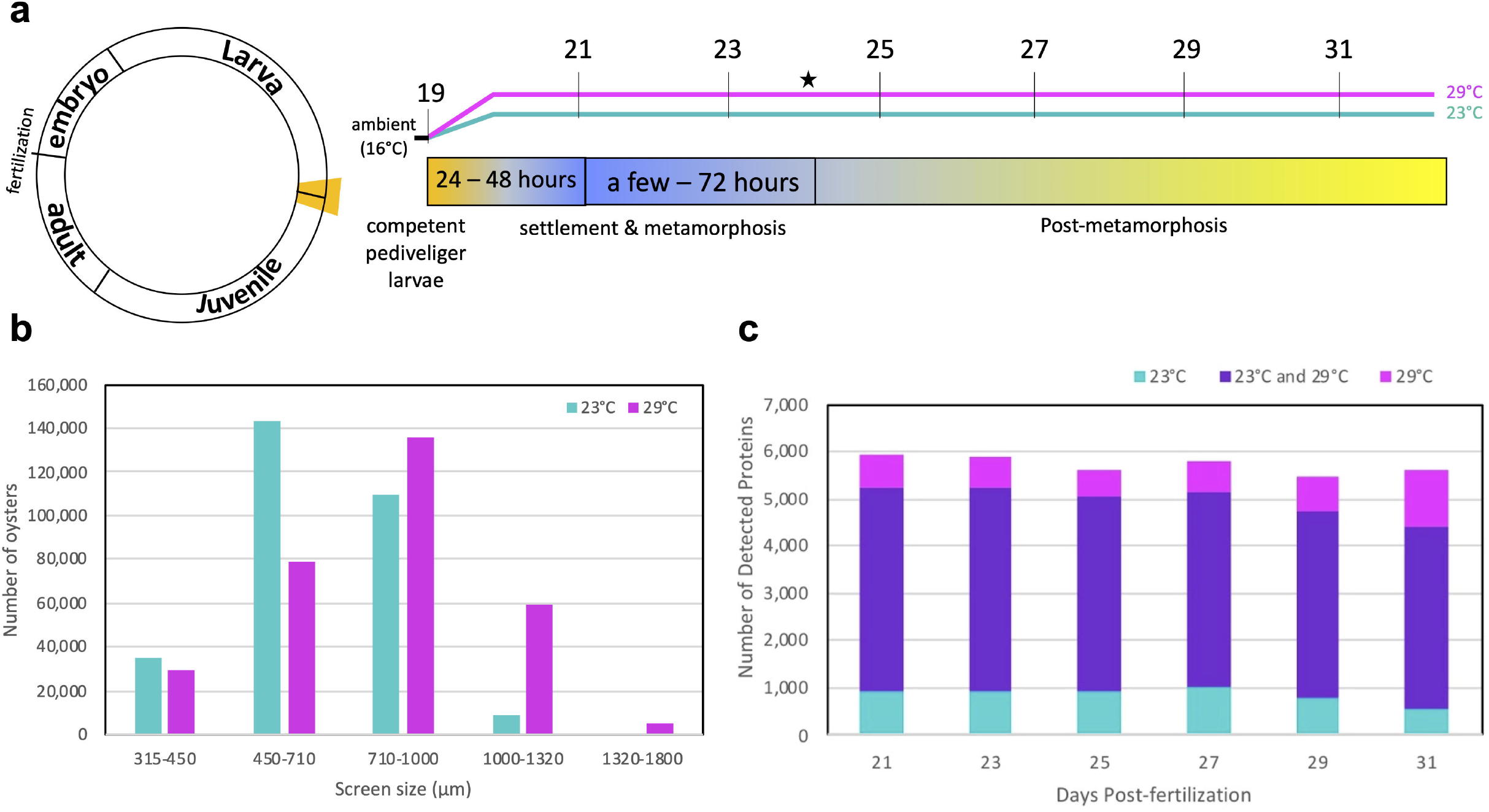
Pacific oyster developmental proteome during metamorphosis. (**a**) Diagram depicting life cycle period examined and the collected time points in days post fertilization at each temperature regime. Color bar shows typical timing of metamorphic transitions. The time point when settlement was assessed is denoted by a star. (**b**) Size distribution based on sorting screen size of oysters at 24 days post-fertilization when settlement was assessed. (**c**) Number of detected proteins at each time point across two rearing temperatures (23°C, cyan; 29°C, magenta, present in both, purple).

Quantitative time-series proteomic measurements resulted in an average of 95,506 ± 2,624 (s.d.) total acquired spectra per sample, with 32.22 ± 2.87% (s.d.) of spectra uniquely mapping to 24,355 ± 2,027 (s.d.) different peptides. All samples collectively covered 19.6% (7,978 out of 40,637 predicted proteins) of the *C.giags* Gigaton database [26], with proteomes at each time point consisting of on average 4,936 ± 255 (s.d.) proteins. Technical replicates from the same time point clustered together and showed less variability than samples from different time points, demonstrating the high reproducibility of our sample preparation and LC MS/MS measurements (**Additional File 2: Supplemental Figure 1**). There was an average overlap of 84 ± 3% of proteins identified in samples from the same time point but different rearing temperature (**Figure 1C**, **Additional File 3: Supplemental Table 2**). A principal components analysis revealed temperature most strongly influenced protein abundance patterns at 21 and 27 dpf, while at 23, 25, 29, and 31 dpf temperature had less of an influence (**Figure 2a)**. From this analysis a total of 70 proteins were identified as top contributors to this temperature-influenced proteomic variation at 21 and 27 dpf by ordering proteins by their greatest magnitude PC 1 and PC2 loadings values and placing a threshold at the first point with the least visual difference between subsequent loadings values (**Additional File 2: Supplemental Figure 2**). These proteins showed three general abundance patterns: decreased abundance at 29°C relative to 23°C at 21 dpf (pale yellow clade), decreased abundance at 29°C relative to 23°C at 27 dpf (salmon clade), and increased abundance at 29°C relative to 23°C at 27 dpf (pale purple clade) (**Figure 2b**, **Additional File 4: Supplemental Table 3**). Proteins with decreased abundance at 29°C relative to 23°C at 21 dpf show significant enrichment of cytoskeleton and extracellular matrix organization, cell motility and locomotion processes (**Figure 2c**). Proteins with decreased abundance at 29°C relative to 23°C at 27 dpf have significant enrichment of early stage development as well as stress response, transport, and catabolic processes, while proteins with increased abundance at 29°C relative to 23°C at 27 dpf have significant enrichment of cellular component and protein complex assembly, transport, and immune system processes. (**Figure 2c**).

**Figure 2.**
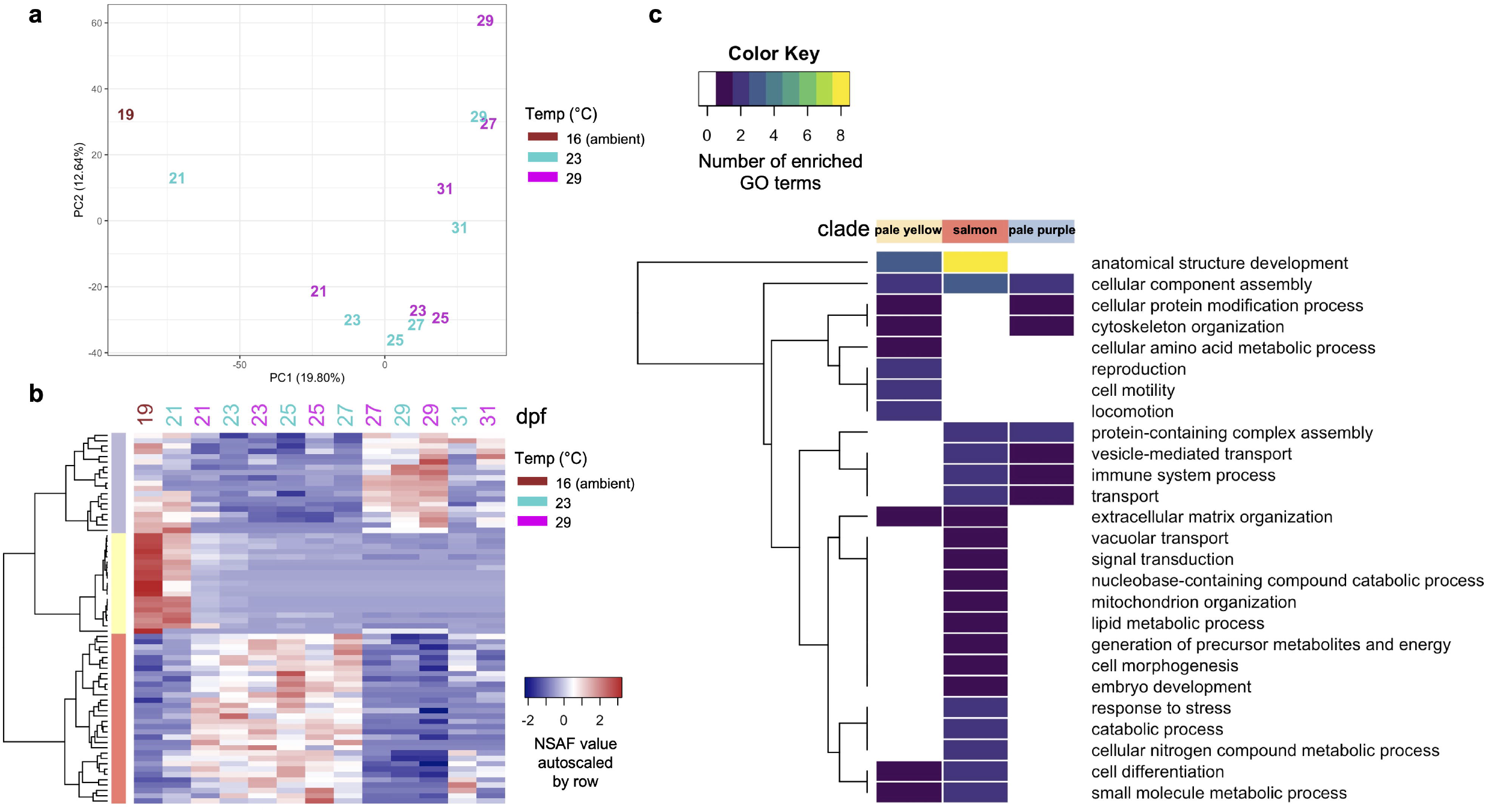
Temperature most influences 21 and 27 dpf proteomes. (**a**) Visualization of the first two principal components from principal component analysis separating samples according to their developmental stage and temperature. Samples are labeled by their sampling time point in days post-fertilization (dpf) with color indicating rearing temperature (16°C, brown; 23°C, cyan; 29°C, magenta). (**b**) Protein abundances (NSAF values autoscaled by row) of proteins most influenced by temperature at 21 and 27 dpf. (**c**) Summary of biological processes represented by enriched GO terms within each clade for temperature-influenced proteins at 21 and 27 dpf.

### Time-influenced proteomic variation

An ANOVA-simultaneous component analysis (ASCA) that partitioned effects from time, temperature, and their interaction revealed that time and the interaction of time and temperature contributed to 91.38% of the variation in protein abundances (**Table 1**). However, a permutation test to quantitatively validate ASCA megavariate effects [27] showed that only time had a significant effect on protein abundances (**Table 1**). In examining the time effect components partitioned by ASCA (**Figure 3a**), we found a total of 217 proteins contributed the most to abundance pattern differences across time when proteins were ordered by their time effect PC1 and PC2 component loadings values and a threshold was placed at the first point with the least visual difference between subsequent loadings values (**Additional File 2: Supplemental Figure 3a-d**). Five distinctive clades were identified through cluster analysis of abundance patterns of these time-influenced proteins (**Figure 3b)**. The identified clades exhibit temporal patterns that appear generally independent of temperature: high abundance early on then acutely reduced (light blue clade), a gradual decrease in abundance over time (purple clade), oscillating abundance that increases through 25 dpf then decreases through 29 dpf and finally increases again at 31dpf (green clade), oscillating abundance that decreases through 25 dpf then increases through 29 dpf and finally decreases at 31dpf (gray clade), and a gradual increase in abundance over time (black clade) (**Figure 3c**).

**Figure 3.**
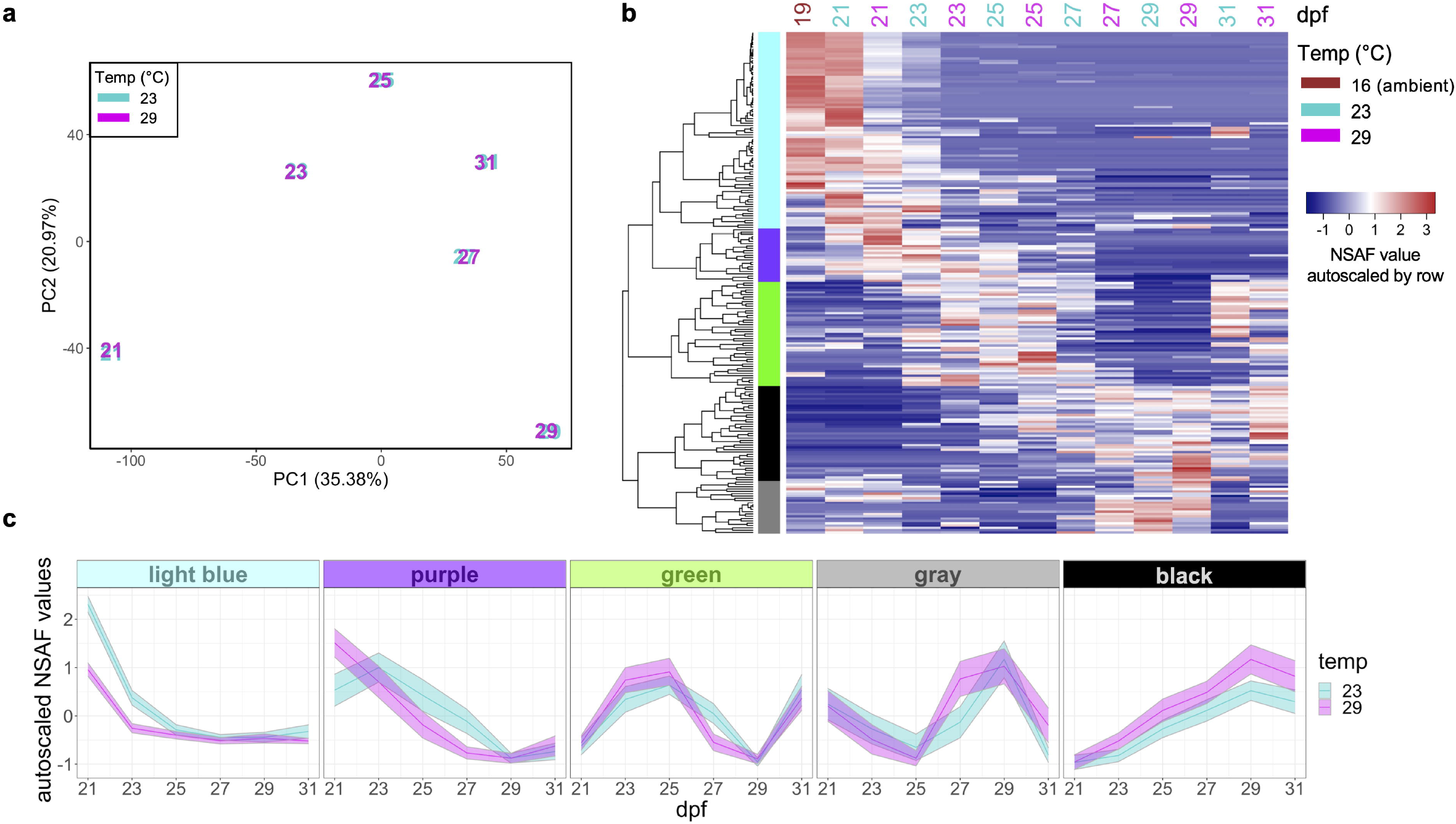
Influence of time on proteomes. (**a**) Score plot for the time factor calculated by ANOVA-simultaneous component analysis of all protein abundances. Samples are labeled by their sampling time point in days postfertilization (dpf) with color indicating rearing temperature (23°C, cyan; 29°C, magenta). (**b**) Protein abundances (NSAF values autoscaled by row) of 217 proteins that contributed the most to time-influenced proteomic variation. (**c**) Temporal abundance patterns of 217 time-influenced proteins based on 5 clades. Bolded line, autoscaled NSAF clade mean with 95% confidence intervals shown.

**Table 1.**
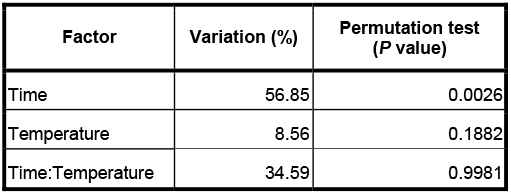
Contributions of experimental factors to the ASCA-partitioned data variation and permutation validation test results.

| Factor | Variation (%) | Permutation test (*P* value) |
| --- | --- | --- |
| Time | 56.85 | 0.0026 |
| Temperature | 8.56 | 0.1882 |
| Time:Temperature | 34.59 | 0.9981 |

Biological processes associated with anatomical structure development and cell differentiation were commonly enriched among proteins across most clades, however, there was a distinct set of enriched biological processes for each clade suggesting specific functions for proteins within each clade (**Figure 4**, **Additional File 5: Supplemental Table 4**). Proteins showing high abundance early on then an acute decrease (light blue clade) were related to immune system, stress response, cell proliferation, cell adhesion, nucleocytoplasmic transport, and cellular amino acid metabolism. Proteins showing a gradual decrease in abundance over time (purple clade) were related to cellular component assembly and protein complex assembly processes. Proteins with oscillating abundance that first increases, then decreases, then increases again (green clade) were related to protein modification, stress response, signal transduction, cell death, and transport biological processes. Different than other clades, this clade had more enriched GO terms related to lipid metabolic process, cell death, cellular amino acid metabolic process, locomotion, and cell motility. Proteins with oscillating abundance that first decreases, then increases, then decreases again (gray clade) were associated with cytoskeleton organization, transport (protein localization, lysozyme transport), and morphogenesis. Different than other clades, this clade had more enriched GO terms related to transport (vesicle transport, vacuolar transport), cell-cell signaling, cytoskeleton organization, protein maturation and developmental maturation. Proteins showing a gradual increase over time (black clade) were largely related to growth and development processes, particularly energy-generating glycolytic processes (e.g. fructose metabolism, mannose metabolism, oligosaccharide metabolism, ganglioside catabolism, and adenosine catabolism) and neurogenesis. These proteins had significant enrichment of GO terms related to carbohydrate metabolic process, cofactor metabolic process, generation of precursor metabolites and energy, nervous system process, plasma membrane organization, and nucleobase-containing compound catabolic process.

**Figure 4.**
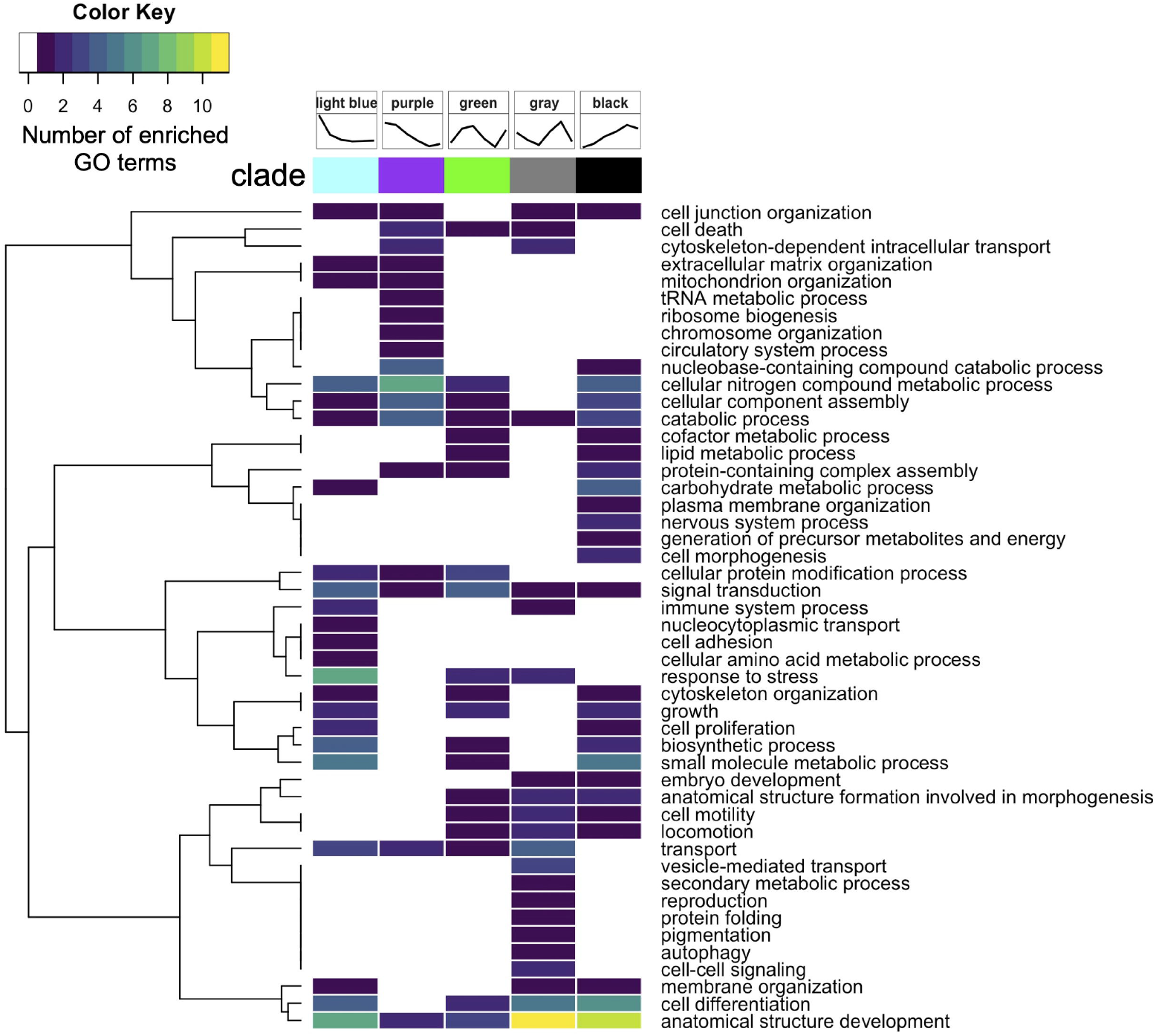
Summary of biological processes represented by enriched gene ontology terms within each clade for time-influenced proteins.

### Proteome response to temperature

Although the ASCA-partitioned effect of temperature was not validated as significant by permutation test (**Table 1**), we still explored proteins contributing to the ASCA-modeled separation of samples by temperature. In examining the temperature effect component partitioned by ASCA (PC1, **Figure 5a**), we found a total of 259 proteins to be significantly influenced by temperature when proteins were ordered by their temperature effect PC1 component loadings values and a threshold was placed at the first point with the least visual difference between subsequent loadings values (**Additional File 2: Supplemental Figure 3e-f**). Hierarchical clustering of these protein abundance patterns revealed two distinctive clades that generally show increased or decreased abundance in 29°C relative to 23°C samples throughout time (orange clade and dark teal clade, respectively; **Figure 5b-c**, **Additional File 6: Supplemental Table 5)**. Proteins showing increased abundance in 29°C relative to 23°C (orange clade) were enriched for growth and development related processes while proteins showing decreased abundance in 29°C relative to 23°C (dark teal clade) were enriched for transport, catabolism, and immune system related processes (**Figure 6**). Immune related proteins in the dark teal clade include putative RNA helicase DEAD box proteins 47 and 58 involved in cellular response to exogenous dsRNA and a putative exosome complex component protein involved in RNA degradation; two homologs of Heme-binding protein 2 and a putative Receptor-interacting serine/threonine-protein kinase 1 known to be involved in positive regulation of necrotic cell death; a putative DnaJ Heat Shock Protein Family (Hsp40) Member A3 known to be involved in cell death activation and growth inhibition; a putative cytidine deaminase known to be involved in signaling and growth inhibition; and a putative COMMD9 involved in neutrophil degranulation.

**Figure 5.**
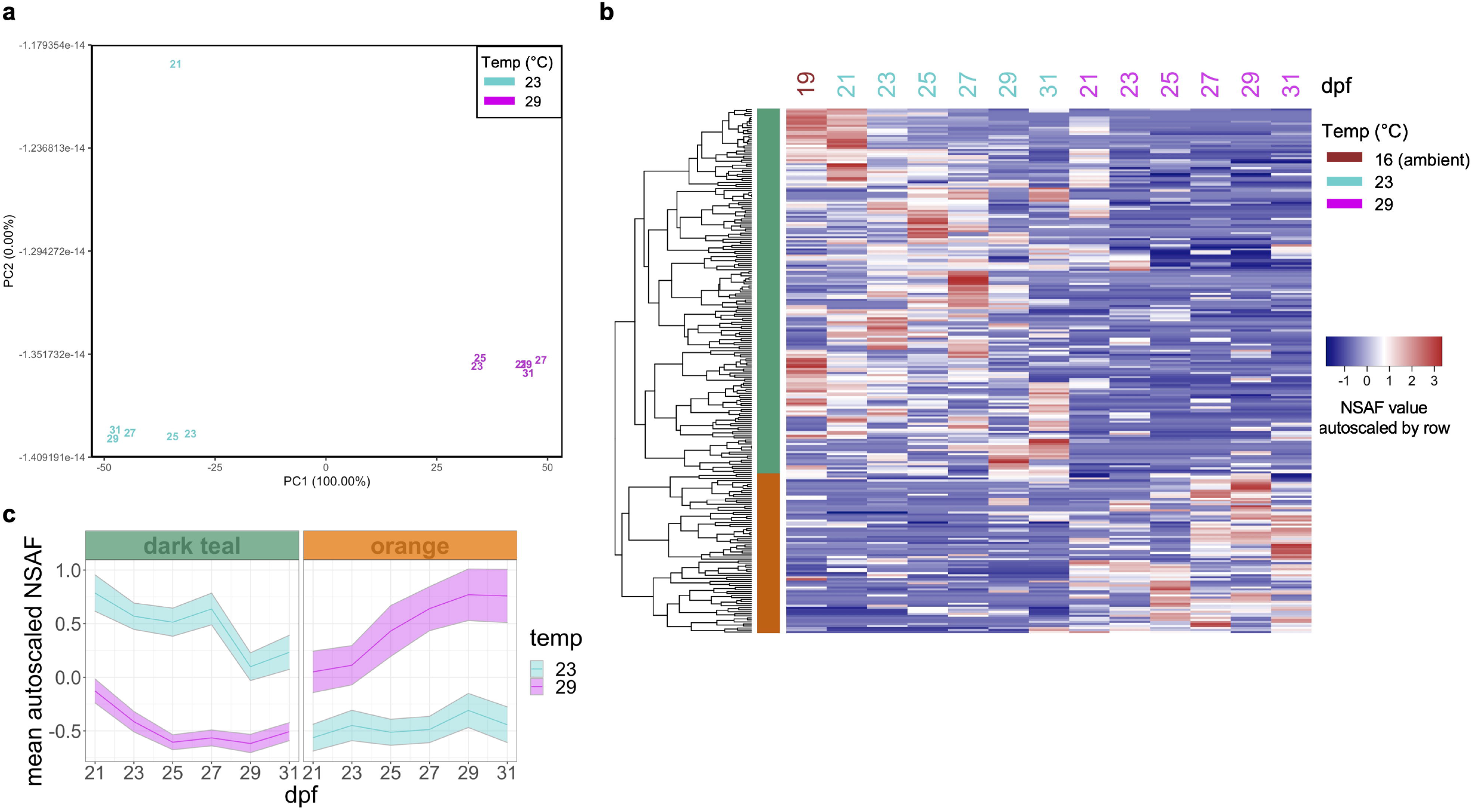
Temperature influence on proteomes. (**a**) Score plot for the temperature factor calculated by ANOVA-simultaneous component analysis of all protein abundances. Samples are labeled by their sampling time point in days post-fertilization (dpf) with color indicating rearing temperature (23°C, cyan; 29°C, magenta). (**b**) Protein abundances (NSAF values autoscaled by row) of 259 proteins that contributed the most to temperature-influenced proteomic variation. (**c**) Temporal abundance patterns of 259 temperature-influenced proteins based on 2 clades. Bolded line, autoscaled NSAF clade mean with 95% confidence intervals shown.

**Figure 6.**
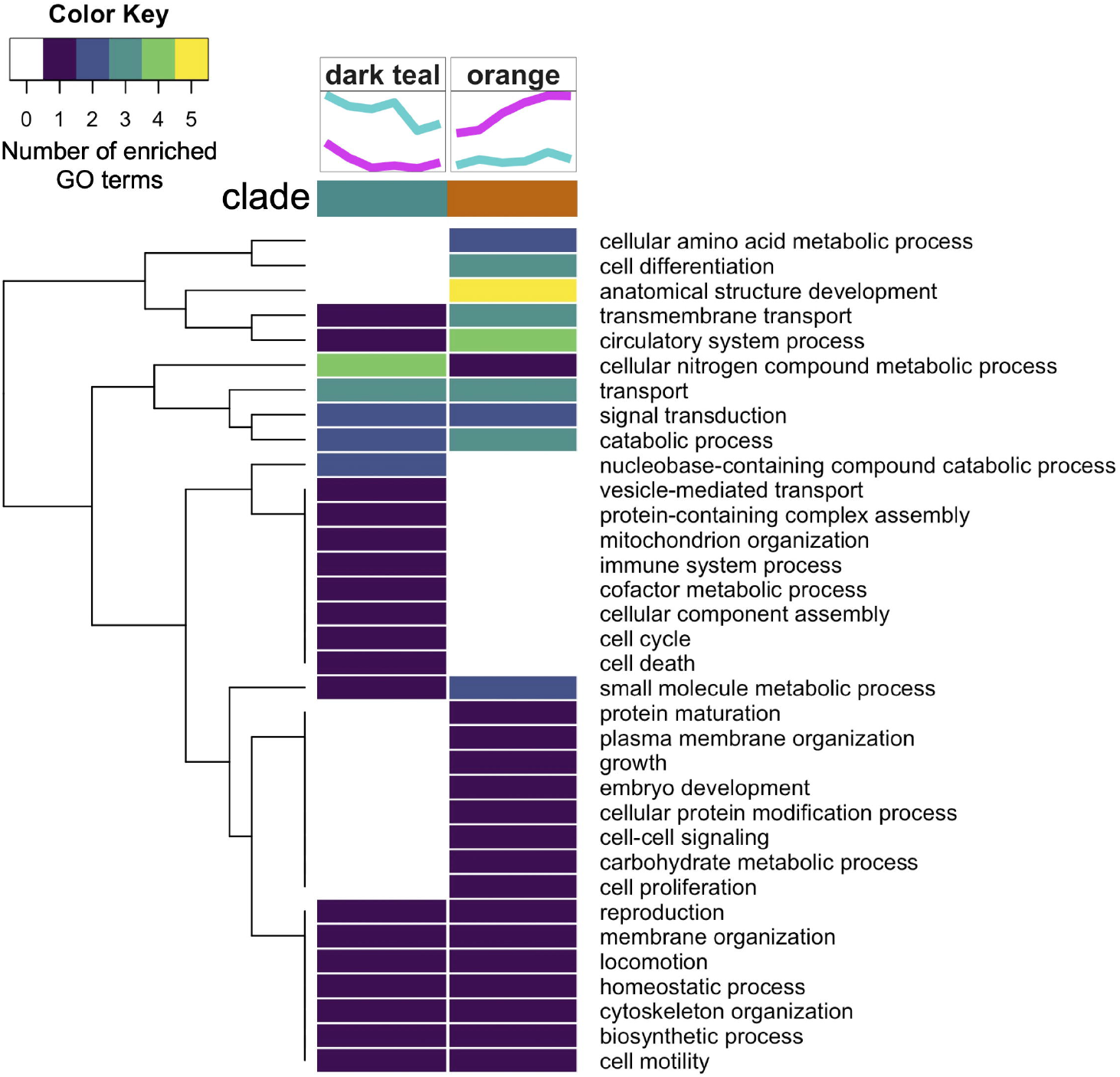
Summary of biological processes represented by enriched GO terms within each clade for temperature-influenced proteins.

## DISCUSSION

We used time series proteomics to explore how oyster physiology is affected by rearing temperature during metamorphosis, generated comprehensive proteomes at seven different time points and across two different temperatures, and did a comparative analysis of how time and temperature affect proteins and their associated biological processes. Performing ASCA, a method which generalizes analysis of variance to a multivariate case [25], allowed the variation in the proteomic data to be partitioned by the experimental factors of temperature, time and their interaction. From this partitioning, we were able to identify proteins that most contribute to variation across time points and variation between temperatures. We were able to further classify differentially abundant proteins by performing hierarchical clustering of their abundance patterns and assigning clades to distinct patterns of abundance change over time and across temperatures. This multi-statistical approach led to the identification of proteins with time and temperature-dependent abundance patterns (**Figures 3** and **5**), and while we could have been more precise in thresholding PC loadings to select proteins, we chose to be more inclusive as to not limit the scope of this exploratory study. Although time and the interaction of time and temperature explained more of the ASCA-partitioned variation in the proteomes, a permutation validation test revealed time had the greatest effect on protein abundance, followed by temperature, and lastly by the interaction of time and temperature (**Table 1**). We chose to not examine the time:temperature interaction effect because the permutation validation test suggested our observations were not different than random chance. However, we chose to examine the temperature effect because it was nearer to being significant and examining proteins contributing to ASCA-partitioned temperature variation allowed two distinct temperature-dependent abundance patterns to emerge. The functional analysis carried out here was limited by few bivalve genes and proteins having been functionally characterized, and mostly depended on gene ontology inferences made from annotated proteins in other species that shared protein sequence similarity to *C. gigas* proteins. Moreover, a number of *C. gigas* proteins with potentially crucial roles in metamorphosis that lacked annotations due to low similarity to proteins in other species in the UniProtKB database [28] were excluded from the functional analysis. Thus, the results from the functional analysis generally should be considered with some uncertainty. Nonetheless, through examining proteins contributing to time and temperature variance we were able to provide comprehensive insight into the proteomic landscape throughout oyster development in conjunction with the nuanced influence of temperature.

We found that temperature only moderately affected proteomes, similar to previous findings [22], and that time was a stronger driver of proteomic differences across samples. This is consistent with the concept that gene expression patterns enable underlying physiological programs to be preserved in different environmental conditions in order for oysters to achieve key developmental stages [22,29,30]. The five distinct protein abundance trends over time are suggestive of particular developmental processes persisting and remaining in sync despite temperature differences.

During typical development, swimming pediveliger larvae settle around 18 dpf, and within 1 to 3 days after settlement most larvae complete metamorphosis to the juvenile stage [5]. In our experiment, two days after larvae appeared competent at 21 dpf, proteins that had high abundance and were then acutely reduced over time (light blue clade) aligned with larval settlement. Seven of the 42 proteins with Uniprot annotations in this clade, and no proteins from other clades, previously showed pediveliger stagespecific expression (CGI_10004853, CGI_10010615, CGI_10006921, CGI_10026725, CGI_10010375, CGI_10006922, and CGI_10006919) [2]. Among other light blue clade proteins were three protease inhibitors, nine structural proteins mostly related to collagen, and five calcification-related proteins that did not previously show pediveliger stage-specific transcript expression, but support hypotheses previously formulated about adhesion during larval settlement [2]. These differences between transcript and protein abundances during settlement highlight the importance of proteomics studies to complement transcriptomics studies.

A key feature of metamorphosis following pediveliger settlement is organ revolution in an anterior-dorsal direction [5]. Proteins that had high abundance at 23 dpf followed by a gradual decrease over time (purple clade) showed clade-specific enrichment of processes related to cell component assembly and protein complex assembly. These included structural proteins (dynein 2 light chain and laminin B2), cell fate determining proproteins (ADAM-TS 16, ADAM-TS 17, and Notch3), and a protein catabolism promoting protein (Cell death regulatory protein GRIM-19), all of which support a tight regulation of growth, cell differentiation and movement that could underlie organ rearrangement at this time point. Moreover, the exosome complex component protein (RRP40) related to the clade-specific enrichment of nucleobase-containing compound catabolism, tRNA metabolic and ribosome biogenesis related processes may play a role in modulating the abundance of pediveliger-specific and recently metamorphosed juvenile-specific lincRNAs [31].

Metamorphosis from larval to juvenile stage encompasses two opposing processes that occur in synchrony: degradation (e.g. velum and foot degradation) and growth processes (e.g. gill and adductor muscle development). We observed two opposing trends in protein abundance patterns over time that were generally unaffected by temperature. Proteins with abundances that increase through 25 dpf then decrease through 29 dpf and increase again at 31dpf (green clade) showed clade-specific enrichment of protein modification and signal transduction related processes generally related to growth promoting pathways. For instance green clade proteins Map kinase kinase 4, Map kinase kinase 5, and PAK-1 are all members of the MAPK signaling pathway that activates cell differentiation and proliferation, potentially related to the development of gill tissue during the prodissoconch and dissoconch postlarval stages and the development of the adductor muscle in the early juvenile stage [5]. Additionally, proteins underlying clade-specific enrichment of lipid and cofactor metabolic processes include oxidative stress protective enzymes paraoxonase and glucose-6-phosphate dehydrogenase which likely act to counter the reactive oxygen species resulting from the aerobic respiration that metamorphosis requires [5]. Proteins with abundances showing an opposing trend decreasing through 25 dpf, then increasing through 29 dpf and decreasing again at 31dpf (gray clade) had clade-specific enrichment of cell-cell signaling and transport processes. These included a protein involved in neurotransmitter release regulation (Snapin), GABA type A receptor-associated protein, Hsc70-interacting protein, and a number of cytoskeleton related proteins (Rho1, Tubulin-folding cofactor B, Filamin-B, and muscle actin LpM) involved in growth regulation. The increased abundance of these proteins at 27-29 dpf could serve in signaling the downregulation of processes initiated by the growth-related proteins at 23-25 dpf (green clade). The potential growth-curbing role of the gray clade proteins is further supported by their decreased abundance at 31 dpf and the enrichment of growth promoting and carbohydrate metabolic processes in proteins showing an increase over time peaking at 31 dpf (black clade). At this time point, the high abundance of fatty acid binding proteins (H-FABP and B-FABP), muscle growth-related proteins (Thymosin beta, Tropomyosin, and Collagen alpha-3 (VI) chain), and carbohydrate metabolic proteins (Hexokinase type 2, Fructose-bisphosphate aldolase, and Ganglioside GM2 activator (involved in the degradation of oligosaccharide-containing gangliosides [32])) is suggestive of muscle tissue building and maintenance (e.g. adductor muscle) that occur after the establishment of structural components, and that the final juvenile stage has been reached.

Although temperature did not have as dominant an effect as time on protein abundances, the two distinct temperature-dependent protein abundance patterns and their associated biological processes support the phenotypic differences we observed in larvae reared at different temperatures. The growth and development-related processes enriched among proteins showing increased abundance in 29°C relative to 23°C regardless of time suggest a mechanism for the larger sizes observed for animals reared 29°C compared to 23°C at 24 dpf. While larger size could be due to elevated temperature increasing overall development rate [33,34] by increasing ingestion activity [13,35], we did not detect a significant time:temperature interaction effect. We did, however, observe specific time points where temperature appeared to have a greater impact on protein abundance (21 and 27 dpf) and thus on timing of key developmental processes. For example, proteins showing increased abundance at 21 dpf (pale yellow clade) related to motility at 23°C (e.g. cilia and flagella associated proteins 53, 54, 58, and 43, TPR repeat protein 25, and tektin-4) could indicate the persistence of the velum organ at 23°C and its degradation at 29°C. The decreased abundance of muscle forming proteins, filament proteins, and signaling proteins at 27 dpf in animals reared at 23°C (pale purple clade) suggests that muscle formation is less robust at 23°C than at 29°C. Of the 79 proteins showing increased abundance regardless of time in 29°C relative to 23°C d c(orange clade), 11 have previously shown increased abundance in response to various 30°C conditions in a proteomics study on pediveliger (15-17 dpf) Pacific oyster [22] (**Table 2**). Nine of these proteins are characterized and are commonly involved in smooth muscle and neuron formation.

**Table 2.** Proteins that commonly show increased abundance in response to high temperature conditions across proteomics studies.

| Protein ID | Uniprot ID | Gene ID | Protein name | Function |
| --- | --- | --- | --- | --- |
| CHOYP_contig_043280.m.49983 | K1PZS2 | CGI_10005951 | Uncharacterized protein | unknown |
| CHOYP_LOC100705966.1.1.m.45957 | K1QJR4 | CGI_10019738 | Heat shock protein beta-1 | stress resistance and actin organization [47], regulates transport of neurofilament proteins [47,48] |
| CHOYP_ADD.3.5.m.17639 | K1PEX5 | CGI_10006848 | Protein hu-li tai shao | actin assembly, important for neuromotor function |
| CHOYP_LOC100367954.2.2.m.66596 | K1P9U4 | CGI_10005881 | Uncharacterized protein | unknown |
| CHOYP_LOC100375029.6.10.m.36981 | K1QM61 | CGI_10009700 | Uncharacterized protein | actin monomer binding [GO:0003785]; actin filament organization [GO:0007015] |
| CHOYP_LOC100375029.8.10.m.60484 | K1QM61 | CGI_10009700 | Uncharacterized protein | actin monomer binding [GO:0003785]; actin filament organization [GO:0007015] |
| CHOYP_LOC101173335.4.4.m.49816 | K1PFT9 | CGI_10006016 | Transgelin | actin cross-linking/gelling protein in fibroblast and smooth muscle tissue [49–51] |
| CHOYP_NF70.1.4.m.31159 | K1PWQ2 | CGI_10018067 | 60 kDa neurofilament protein | intermediate filament [GO:0005882] |
| CHOYP_RPS24.1.8.m.571 | K1PUV4 | CGI_10001493 | 40S ribosomal protein S24 | ribosome [GO:0005840]; structural constituent of ribosome [GO:0003735]; translation [GO:0006412] |
| CHOYP_contig_044078.m.50900 | K1Q086 | CGI_10019530 | Ankyrin-2 | essential role in localization and membrane stabilization of ion transporters in muscle cells [52,53], signal transduction [GO:0007165] |
| CHOYP_LOC100696604.1.1.m.40638 | K1QP17 | CGI_10010975 | Caprin-1 | directly binds mRNA involved in neuronal synaptic plasticity, cell proliferation, and migration in multiple cell types; may regulate mRNA transport and translation [54] |

The immune system, vesicle transport, and nucleobase-containing, nitrogen compound catabolic processes associated with proteins showing decreased abundance in animals reared at 29°C (dark teal clade) are likely associated with metamorphic differences (e.g. different degradation rates of velum or foot), differences in metabolic rate, differences in pathogen presence or susceptibility, or a combination of these. Seawater pathogen presence and abundance could have been different between temperature treatments since temperature can alter seawater microbiome composition [36]. For instance, ciliates from the genus Orchitophryiadae that infect marine invertebrates [37] have shown an upper thermotolerance limit of 27°C [38], and might not have survived at 29°C but could have at 23°C. Moreover, temperature can affect both pathogenicity and host susceptibility. For instance, Pacific oyster juveniles infected with OSHV-1 had significantly higher survival at 29°C than 21°C, potentially because at 29°C they are able to alter their physiology to limit viral entry [39,40]. Specifically related to the proteomics results, viral recognition protein DEAD box protein 58 and growth inhibition/cell death-related proteins serine-threonine kinase receptor-associated protein and DnaJ-like protein have been implicated in dsRNA exposure response in adult Pacific oyster gill tissue [41]. Larval geoduck proteomic response to ciliates similarly showed an increase in immune response proteins including molecular chaperones and reactive oxygen species responders [42]. Reduced abundance of these proteins in animals at 29°C suggests they were not sustaining immune and cytoprotection processes, allowing them to reallocate energy to growth [43]. Moreover, potential pathogen exposure in animals reared at 23°C could have led to a decline in feeding activity and therefore less growth [43]. A delay in velum degradation in oysters reared at 23°C could have led to an immune challenge as the velum in particular has shown susceptibility to infection [44–46]. Temperature influenced proteomic differences could therefore be confounded by differences in seawater microbial composition. With no obvious source of pathogen exposure, further controlled trials are needed to determine the extent to which these specific proteome differences are driven by pathogen presence and abundance differences, metabolic rate differences, and/or specific metamorphic process differences.

## CONCLUSION

We successfully simultaneously surveyed thousands of proteins to generate comprehensive proteomes from which we were able to identify temporal and temperature-influenced protein abundance patterns. We characterized physiological processes related to proteins showing different abundance patterns that underlie core developmental processes, and that underlie the effects of different temperature regimes. These findings offer high resolution insight into the role of temperature and why oysters may experience high mortality rates during this life transition in both field and culture settings. The proteome resource generated and the temporally mapped physiological processes offer data-driven guidance for further investigating specific metamorphic stage transitions, temperature-related phenotypic differences in mollusc development, and developmental regulation in general. Lastly, the analytical approach taken here provides a foundation for effective shotgun proteomic analyses across a variety of taxa.

## METHODS

### Larval rearing

All seawater used for rearing flowed continuously from Dabob Bay, WA at 3 L/min, filtered to 5μm, and was maintained at pH 8.4 by the addition of sodium carbonate. Animals were fed *T-Isochrysis, Pavlova* sp., *Nannochloropsis* sp., *Rhodomonas* sp., and *Tetraselmis* sp. at constant effluent algal densities of 100,000 cells/mL throughout the experiment. *Crassostrea gigas* (triploid) larvae were reared at ambient temperature (16°C) until they developed eyespots and pedal appendages, and were greater than 250 μm in size (pediveliger stage; 19 days post-fertilization). An initial sample of ~12,500 pediveliger larvae was rinsed with filtered seawater, dried, flash frozen in liquid nitrogen and stored at −80°C for proteomics analysis. Approximately one million pediveligers (20.0 g) were transferred to the experimental rearing system consisting of a silo (46 cm PVC pipe) containing 80mL of ground oyster shell (180-315 μm graded microculch) as settlement substrate. Each silo was housed inside a rearing tank with flowing filtered seawater and held at either 23°C or 29°C for 13 days. Tanks were drained and oysters were rinsed with filtered seawater daily.

### Settlement assessment and sampling

At 24 days post-fertilization, settlement and size were assessed using sorting screens ranging from 450 to 1320 μm, and any larvae that had not set were removed. Settlement was calculated as the proportion of larvae captured on the sorting screens. At 21, 23, 25, 27, 29, and 31 days post-fertilization (corresponding to days 3, 5, 7, 9, 11, and 13 of the experiment, respectively) samples of approximately 12,500 larvae were collected from each of the two temperature regimes (23°C or 29°C) for shotgun proteomics analysis. Larvae were rinsed with filtered seawater, dried, flash frozen in liquid nitrogen and stored at −80°C. In total there were 13 samples taken for proteomic analysis, one initial sample from 19 days post-fertilization at ambient temperature (16°C) and six at each temperature regime throughout development.

### Protein Sample Preparation

Cell homogenates were prepared by adding 500 μL of 50 mM NH_4_HCO_3_ in 6 M urea to the sample and homogenizing with a pestle directly in the microfuge tube. Samples were centrifuged at 2000 rpm for 5 minutes to separate solid shell and tissue fragments from the cellular content fraction (the supernatant), and the supernatant (150 μL) was transferred into new tubes. Supernatants were sonicated three times each for 5 seconds, cooling samples in between sonication rounds using an ethanol and dry ice bath for 5 seconds. After sonication, sample protein concentrations were determined using a BCA assay kit (Pierce). Protein digestion was carried out by diluting 100 μg of protein from each sample with 50 mM NH_4_HCO_3_ in 6 M urea solution to a final volume of 100 μL, adding 1.5 M tris pH 8.8 (6.6 μL) and 200 mM tris (2-carboxyethyl)phosphine hydrochloride (2.5 μL) and vortexing samples. Samples were maintained at a basic pH > 7 by titrating with sodium hydroxide (5N). After incubating samples for one hour at 37°C, 20 μL of 200 mM iodoacetamide was added, samples were vortexed then incubated for one hour at room temperature in the dark. Next, 20 μL of 200 mM diothiothreitol was added, samples were vortexed, and incubated for one hour at room temperature. Then, 1.65 μL LysC (lysyl endopeptidase, Wako) (1:30 enzyme:protein ratio) was added to each sample, samples were vortexed, and incubated for one hour at room temperature. Finally, 800 μL 25 mM NH_4_HCO_3_ 200 μL HPLC grade methanol and 3.3 uL Trypsin (Promega) (1:30 enzyme:protein ratio) were added to each sample, samples were vortexed, and incubated overnight at room temperature. Samples were evaporated using a centrifugal evaporator at 4°C to near dryness and stored at −80°C. Desalting of samples was done using Macrospin columns (sample capacity 0.03-300 ug; The Nest Group, Southborough, MA) following the manufacturer’s instructions. Dried peptides were reconstituted in 100 μL 3% acetonitrile + 0.1% formic acid and stored at −80°C.

### Mass Spectrometry

Data-dependent acquisition was performed on an Orbitrap Fusion Lumos Mass Spectrometer (Thermo Scientific) at the University of Washington Proteomics Resource to assess the effect of temperature on proteomic profiles throughout larval development. Technical duplicates for each sample were processed by liquid chromatography coupled to tandem mass spectrometry (LC-MS/MS). Briefly, the analytical column (20 cm long) was packed in house with C18 beads (Dr. Maisch HPLC, Germany, 0.3 μm) with a flow rate of 0.3 μL/min. Chromatography was carried out with an increasing ratio of acetonitrile + 0.1% formic acid (solvent A):water + 0.1% formic acid (solvent B). The solvent gradient was 5-95% solvent A over 70 min. Quality-control standards (Pierce Peptide Retention Time Calibration mixture + bovine serum albumin) were analyzed throughout the experiment to ensure consistency of peptide detection and elution times.

### Protein identification and quantification

Mass spectrometer raw files (PRIDE Accession no. PXD013262) were converted to .mzXML format and were searched against a protein sequence database that contained the *C. gigas* proteome (downloaded from http://gigaton.sigenae.org [26]) and common contaminants (downloaded from the crapOME [55]) using Comet v. 2016.01 rev.2 [56]. The Trans Proteomic Pipeline [57] was then used to calculate statistics associated with peptide-to-protein matches with a peptide probability *p*-value threshold of 0.9. Next, Abacus [58] was used to correlate protein inferences across samples and obtain a single protein identification for each peptide. From the Abacus output file, the adjusted normalized spectral abundance factor (NSAF) values (spectral abundance normalized to protein sequence length) were used to compare technical duplicates and biological samples in a principal component analysis (PCA) (**Additional File 2: Supplemental Figure 1**). For the PCA, NSAF values were log2 transformed after converting zero values to 0.1 (1/8 of the lowest NSAF value). NSAF values from technical replicates were averaged for each protein in all downstream analyses.

### Preliminary principal component analysis

For a preliminary assessment of the overall variability of proteomes, PCA was run on log2 transformed NSAF values for all samples where NSAF values of zero were converted to 0.1 (1/8 of the lowest NSAF value) prior to log transforming. To identify proteins that most contribute to PC1 and PC2 variation, PC1 and PC2 protein loadings values were ordered and plotted by greatest magnitude loadings value. For each loadings plot, a threshold was placed at the point of diminishing returns, at the first point with the least visual difference between subsequent loadings values, which for PC1 was a loadings value greater than 0.0236 or less than −0.02355, and for PC2 was a loadings value greater than 0.0245 or less than −0.0262 (**Additional File 2: Supplemental Figure 2**). Proteins with a loadings value magnitude greater than or equal to the thresholds were considered proteins most contributing to variation accounting for temperature differences between samples from 21 and 27 dpf.

### ANOVA-simultaneous component analysis

ANOVA-simultaneous component analysis (ASCA) from the R package MetStat [59] was used to evaluate the effect of time, temperature, and their interaction on protein abundance using the log2 transformed average NSAF values for all samples. To quantitatively validate the significance of effects estimated by ASCA, a permutation test was performed that randomly reassigned group labels and recalculated the ASCA sum of squares 10,000 times [27]. To identify proteins influenced by time, PC1 and PC2 loadings values of the ASCA-generated PCA for the time factor were ordered by decreasing magnitude and plotted. For each plot a loadings value threshold was placed at the point of diminishing returns, thresholding at the first point with the least visual difference between subsequent loadings values, which for both PC1 and PC2 was ≥ 0.035 or ≤ −0.035 (**Additional File 2: Supplemental Figure 3a-d**). To identify proteins influenced by temperature, PC1 loadings values of the ASCA-generated PCA for the temperature factor were ordered by decreasing magnitude, plotted, and a threshold was placed at the point of diminishing returns, at the first point with the least visual difference between subsequent loadings values, at ≥ 0.03 or ≤ −0.025 (**Additional File 2: Supplemental Figure 3e-f**).

### Cluster analysis

Hierarchical clustering was performed on temperature-influenced (identified by PCA or ASCA) and time-influenced proteins (identified by ASCA only) using the complete linkage clustering algorithm and Pearson correlation-based distances. Dendrograms generated were cut at the height of 1 for proteins identified as temperature-influenced by PCA, 1.8 for time-influenced proteins identified by ASCA, and 1.5 for temperature-influenced proteins identified by ASCA to define clades within each group of proteins. Heatmaps of temperature- and time-influenced proteins with clades shown as a colored sidebar were generated using the R package Heatmap3 [60] and protein abundance plots were generated using the R package ggplot2 [61].

### Functional enrichment

To explore the biological processes related to proteins influenced by time and temperature, we performed Gene Ontology (GO) enrichment analysis on the proteins within each clade assigned by the aforementioned cluster analysis. First, to retrieve GO annotations protein sequences from the *C. gigas* proteome (downloaded from http://gigaton.sigenae.org [26]) were queried against the UniProt protein database (UniProt release 2019_01 [28]), a comprehensive reference set of protein sequences and functional information including GO annotations from thousands of species using BLASTp [62]. The rationale behind using a multi-species reference set as opposed to a species-specific reference set was to obtain functional information for as many proteins as possible where many *C. gigas* proteins lack annotation and homologs in other species may have annotations. The alignment with the lowest e-value was kept for each protein sequence, and alignments were filtered further for those with an e-value of less than or equal to 1 x 10^-10^ to keep only high confidence alignments. GO annotations from the Uniprot alignments were used for GO enrichment analysis. A Fisher test was performed with TopGO [63] using default settings on proteins from each clade defined by methods described above, using all detected proteins across all samples with GO annotations in the Uniprot alignments as the background set. GO terms with *P* < 0.05 and occuring ≥ 5 times in the background set were considered significant [64]. Correction for multiple testing was not applied on the resulting *P* values because the tests were not considered to be independent [65]. Significant GO terms were converted to GO Slim terms using the R package GSEAbase [66], and heatmaps were generated using the heatmap.2 function in the R package gplots [67].

### Comparison to published datasets

Proteins influenced by time and temperature were compared to genes and proteins previously identified in other *C. gigas* proteomics studies exploring thermal impact on pediveliger stage [22] and an *in silico* transcriptomic study of larval settlement [2] using their Uniprot identifiers. Because the Uniprot identifiers in these published datasets were from only the *C. gigas* species, *C. gigas* Uniprot identifiers were retrieved for temperature- and time-influenced proteins by aligning their protein sequences from the *C. gigas* proteome (downloaded from http://gigaton.sigenae.org [26]) to the species-specific *C. gigas* reference proteome from Uniprot database (‘UP000005408_29159’) using BLASTp, keeping only the alignment with the lowest e-value for each protein sequence. Alignments were filtered for those with an e-value of less than or equal to 1 x 10^-10^ to keep only high confidence alignments. For temperature-influenced proteins that previously were identified and showed increased abundance in pediveliger proteomes arising from various heat exposure conditions [22], functional information was retrieved from querying the Uniprot knowledgebase [28] with their Uniprot identifiers using the Retrieve/ID mapping tool.

## ABBREVIATIONS

ADAM-TS: A disintegrin and metalloproteinase with thrombospondin motifs
ANOVA: Analysis of variance
ASCA: ANOVA-simultaneous component analysis
BCA: Bicinchoninic acid assay
COMMD9: Copper metabolism MURR1 domain-containing protein 9
DEAD box: family of proteins containing a motif with the amino acid sequence asp-glu-ala-asp
dpf: day post-fertilization
dsRNA: double stranded ribonucleic acid
FABP: Fatty acid-binding protein
GABA: Gamma aminobutyric acid
GO: Gene ontology
GRIM-19: Gene associated with retinoid-IFN-induced mortality
Hcs70: Heat shock cognate protein 70
LC-MS/MS: liquid chromatography coupled tandem mass spectrometry
lincRNA: Long intergenic noncoding RNA
MAPK: Mitogen-activated protein kinase
NSAF: Normalized spectral abundance factor
PAK1: p21-activating kinase 1
PCA: Principal component analysis
PVC: polyvinyl chloride
RNA: ribonucleic acid
RRP40: Ribosomal RNA-processing protein 40
TPR repeat: tetratricopeptide repeat
tRNA: transfer ribonucleic acid

## DECLARATIONS

### Ethics approval and consent to participate

Not applicable.

### Consent for publication

This manuscript does not include any personal communications and did not require permission from individuals other than the authors for the use of any unpublished data.

### Availability of data and materials

Mass spectrometry proteomics data (.raw, .pepxml, and .mzXML files) have been deposited to the ProteomeXchange Consortium via the PRIDE (PubMed ID: 30395289) partner repository with the dataset identifier PXD013262. Additional files, additional supporting materials, and code is available at 10.6084/m9.figshare.12436598.

### Competing interests

The authors declare that they have no competing interests.

### Funding

This material is based upon work supported by the Washington Sea Grant award NA140AR4170078; project R/SFA-8.

### Authors’ contributions

S.B.R., B.V., and B.E. conceived the project. S.B.R., B.V., and B.E. advised research. R.E.T. and E.B.T.S. performed experiments. R.E.T. and S.B.R. performed proteomics analysis. S.A.T. performed statistical analyses with help from K.R.M. S.A.T. performed functional analyses. S.A.T. prepared the manuscript with edits from S.B.R., B.V., K.R.M., and E.B.T.S.

## Acknowledgements

We thank the Taylor Shellfish Hatchery for providing the animals, water treatment, rearing systems, and husbandry. We also thank the current and former members of the Roberts lab at the University of Washington as well as the University of Washington’s Proteomics Resource (UWPR95794).

## ADDITIONAL FILES

| **File name** | **File format** | **Title of data** | **Description of data** |
| --- | --- | --- | --- |
| Additional file 1:<br>Supplemental_Table_1 | xlsx | Biovolume of oyster seed (mL) settled after 6 days of exposure to high temperatures | Data and plot of oyster counts and size measurements for Figure 1b |
| Additional file 2:<br>Supplemental_Figures | pdf | PCA of all technical replicate samples | Supplemental Figure 1. PCA of all technical replicate samples<br>Supplemental Figure 2. Preliminary principal component analysis plots of PC loadings for the top 100 ranked proteins.<br>Supplemental Figure 3. ANOVA-simultaneous component analysis plots of PC loadings for all proteins. |
| Additional file 3:<br>Supplemental_Table_2 | xlsx | Preliminary proteome characterization | Data and plot showing overlap between different temperature proteomes at each timepoint for Figure 1c |
| Additional file 4:<br>Supplemental_Table_3 | csv | PCA, hierarchical clustering, and functional analysis results for all detected proteins. | PCA, hierarchical clustering, and functional analysis results for all detected proteins. |
| Additional file 5:<br>Supplemental_Table_4 | csv | ASCA time effect analysis, hierarchical clustering, and functional analysis results for all detected proteins. | ASCA time effect analysis, hierarchical clustering, and functional analysis results for all detected proteins. |
| Additional file 6:<br>Supplemental_Table_5 | csv | ASCA temperature effect analysis, hierarchical clustering, and functional analysis results for all detected proteins. | ASCA temperature effect analysis, hierarchical clustering, and functional analysis results for all detected proteins. |

## Notes

https://doi.org/10.6084/m9.figshare.12436598.v5

### Competing Interest Statement

The authors have declared no competing interest.

### Summary of Updates

Abstract, Figures, Manuscript text, and Supplemental material to address Reviewers' comments

