## Supplemental Figures for "Temporal proteomic profiling reveals insight into critical developmental processes and temperature-influenced physiological response differences in a bivalve mollusc"

**Supplemental Figure 1.** PCA of all technical replicate samples. (page 2)

**Supplemental Figure 2.** Principal component analysis plots of PC loadings for the top 100 ranked proteins. (page 2)

**Supplemental Figure 3.** ANOVA-simultaneous component analysis plots of PC loadings for all proteins. (page 3)

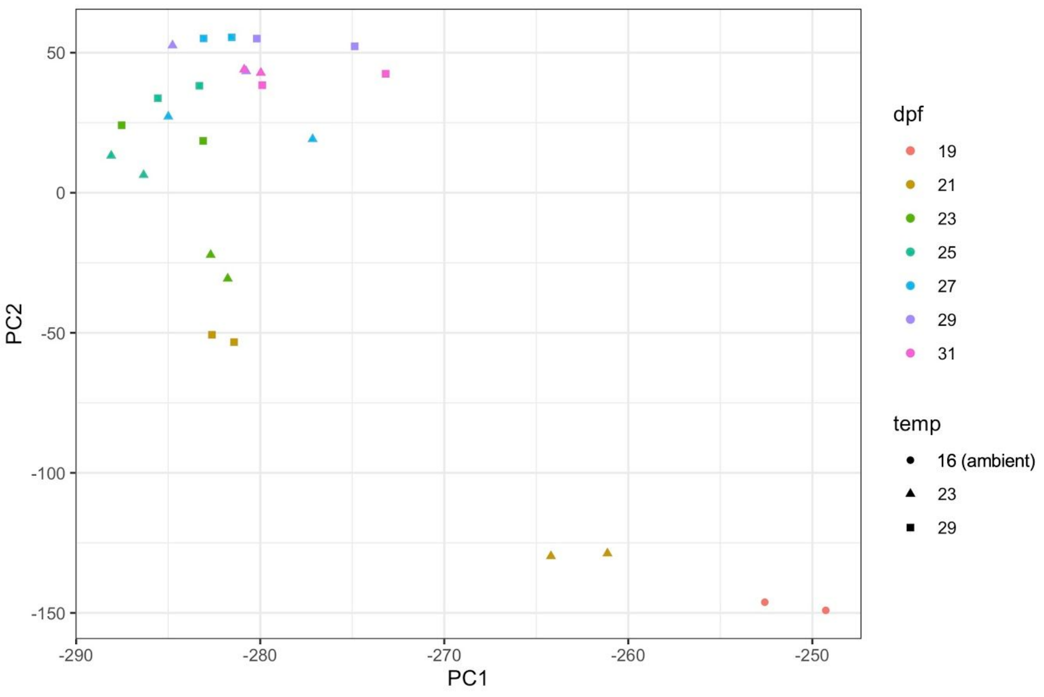


**Supplemental Figure 1.** PCA of all technical replicate samples.

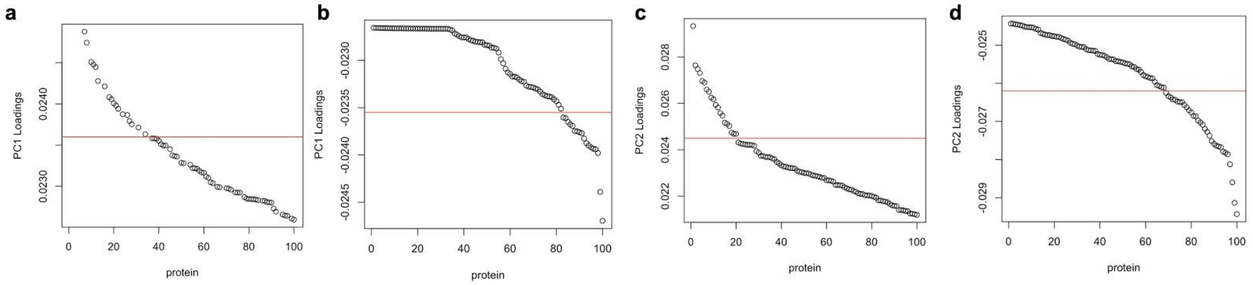


**Supplemental Figure 2.** Preliminary principal component analysis plots of PC loadings for the top 100 ranked proteins. (**a**) Proteins with the top 100 highest positive PC1 loadings thresholded (red line) at 0.0236. (**b**) Proteins with the top 100 lowest negative PC1 loadings thresholded (red line) at -0.02355. (**c**) Proteins with the top 100 highest positive PC2 loadings thresholded (red line) at 0.0245. (**d**) Proteins with the top 100 lowest negative PC2 loadings thresholded (red line) at -0.0262.

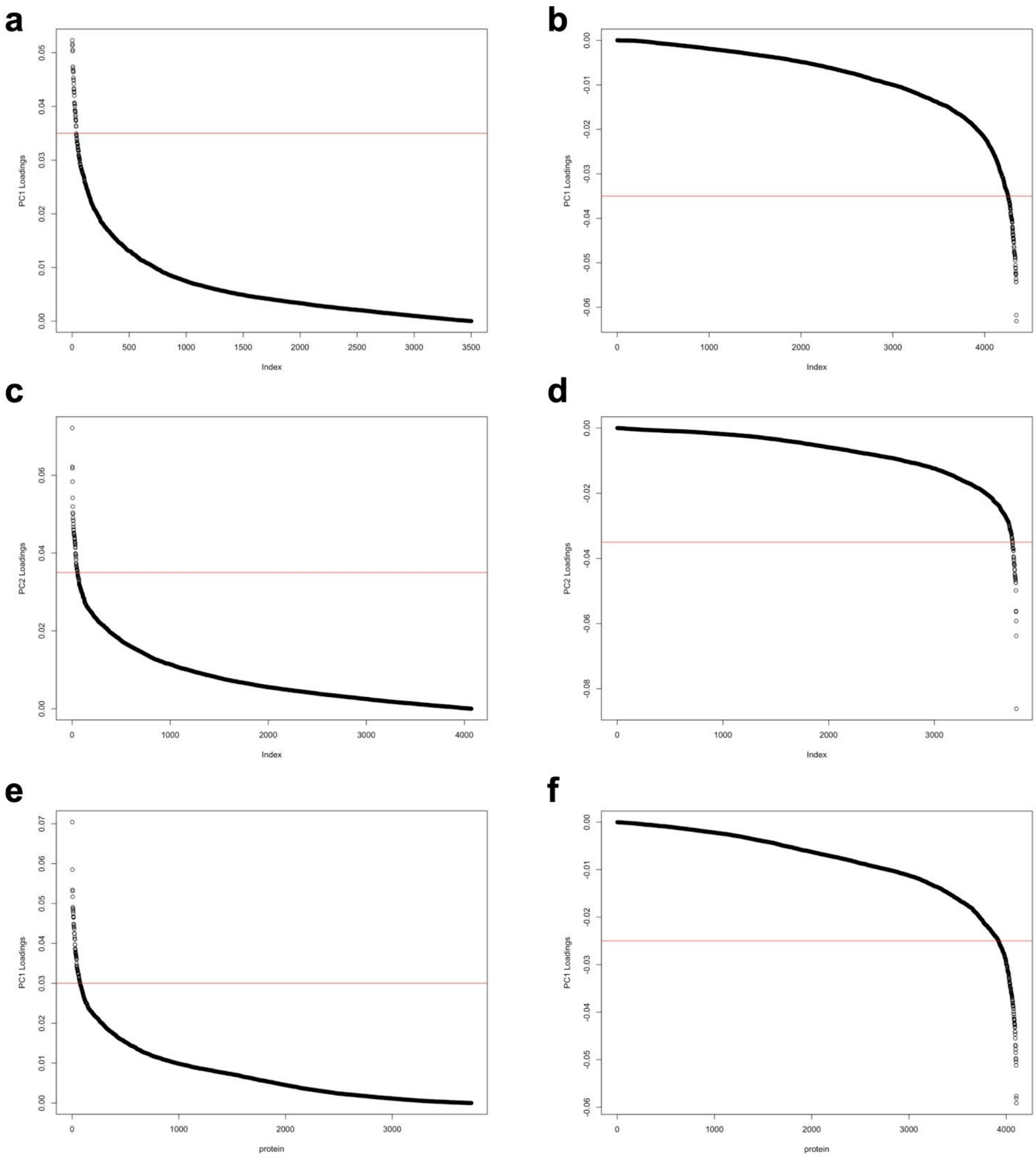


**Supplemental Figure 3.** ANOVA-simultaneous component analysis plots of PC loadings for all proteins. **(a)** Positive PC1 loadings for time effect. (**b**) Negative PC1 loadings for time effect. **(c)** Positive PC2 loadings for time effect. (**d**) Negative PC2 loadings for time effect. (**e**) Positive PC1 loadings for temperature effect. (**f**) Negative PC1 loadings for temperature effect. Red line indicates the significance threshold loadings value set at the point of diminishing returns (0.035, -0.035, 0.035, -0.035, 0.03, and -0.025, respectively).
